# End-product inhibition of the LRRK2-counteracting PPM1H phosphatase

**DOI:** 10.1101/2025.05.16.654599

**Authors:** Ayan Adhikari, Aashutosh Tripathi, Claire Y. Chiang, Pemba Sherpa, Suzanne R. Pfeffer

**Affiliations:** Department of Biochemistry, Stanford University School of Medicine, Stanford, CA, United States; Aligning Science Across Parkinson’s (ASAP) Collaborative Research Network, San Francisco, CA, United States

**Keywords:** Rab, phosphatase, leucine-rich repeat kinase 2 (LRRK2), enzyme inhibitor, Parkinson disease

## Abstract

PPM1H phosphatase reverses Parkinson’s disease-associated, Leucine Rich Repeat Kinase 2-mediated, Rab GTPase phosphorylation. We showed previously that PPM1H relies on an N-terminal amphipathic helix for Golgi membrane localization and this helix enables PPM1H to associate with liposomes *in vitro*; binding to highly curved liposomes activates PPM1H’s phosphatase activity. We show here that PPM1H also contains an allosteric binding site for its non-phosphorylated reaction products, Rab8A and Rab10. Microscale thermophoresis revealed that PPM1H binds thio-phosphorylated Rab8A at the active site with a KD of ∼1µM; binding of Rab8A and Rab10 to an alternative site is of similar affinity and is not detected for another LRRK2 substrate, Rab12. Non-phosphorylated Rab8A or Rab10 inhibit PPM1H phosphatase reactions at concentrations consistent with their measured binding affinities and fail to inhibit PPM1H L66R phosphatase reactions. Independent confirmation of non-phosphorylated Rab binding to PPM1H was obtained by sucrose gradient co-flotation of non-phosphorylated Rabs with liposome-bound PPM1H. Finally, Rab8A or Rab10 binding also requires PPM1H’s amphipathic helix, without which the interaction affinity is decreased about 6-fold. These experiments indicate that Golgi associated Rab proteins contribute to the localization of PPM1H and non-phosphorylated Rabs regulate PPM1H phosphatase activity via an allosteric site. Targeting this site could represent a strategy to enhance PPM1H-mediated dephosphorylation of LRRK2 substrates, offering a potential therapeutic approach to counteract LRRK2-driven Parkinson’s disease.

## Introduction

Activating mutations in Leucine Rich Repeat Kinase 2 (LRRK2) are a major cause of inherited Parkinson’s disease (1, 2). LRRK2 phosphorylates a subset of the 65 human Rab GTPases at a critical Threonine or Serine residue within the Rab’s so-called Switch 2 motifs (3, 4). Rab8A, Rab10 and Rab12 are broadly expressed and prominent LRRK2 substrates, while Rabs 3, 29, 35 and 43 show more tissue specific expression and appear to be phosphorylated at lower stoichiometries (5). It is important to note that at steady state, only a few percent of even the most abundant Rab substrates are phosphorylated (5), and the phosphate modification shows a high rate of turnover, in some cases, with a half-life of as little as 1-2 minutes (6, 7).

Rab phosphorylation blocks the ability of Rabs to bind their normal effector, partner proteins, to be activated by cognate guanine nucleotide exchange factors, and to be retrieved from membranes by GDI proteins (3, 4). Thus, phosphorylation inhibits normal Rab function. However, once phosphorylated, Rabs gain the capacity to bind to phosphoRab-specific effectors, with dominant effects leading to cellular pathology. PhosphoRab effectors identified to date include: RILPL1, RILPL2, JIP3, JIP4, and MyoVa proteins (3, 8, 9).

Berndsen et al. (6) discovered PPM1H as a Rab-specific phosphatase that can reverse LRRK2-mediated Rab phosphorylation. Multiple lines of evidence confirm PPM1H’s role in phosphoRab biology. Loss of PPM1H phenocopies hyperactivation of LRRK2 in cell culture and mouse brain. Upon addition of LRRK2 inhibitors, cells lacking PPM1H display a threefold higher basal level of Rab8 and Rab10 phosphorylation than control cells. Finally, unbiased mass spectrometry of proteins bound to a substrate-trapping PPM1H mutant showed strong enrichment for Rab10, Rab8A, and Rab35 (6).

PPM1H relies upon an N-terminal, amphipathic helix for its membrane association in cells (10). In addition, using purified proteins, we showed that the amphipathic helix can mediate liposome association and confers the capacity of PPM1H to be activated by highly curved membranes (10). In cells, PPM1H is localized near its pRab8A and pRab10 substrates (6, 10) and it can act on pRab12 if the two proteins are artificially co-localized on mitochondrial surfaces (10, 11). These experiments highlight the importance of enzyme and substrate co-localization contributing to apparent substrate specificity in cellular assays.

In our initial, *in vitro* PPM1H-catalyzed Rab de-phosphorylation assays, we noticed that the reactions did not go to completion. Troubleshooting revealed that our phosphoRab10 substrate preparations were not 100% phosphorylated and this led to our discovery that PPM1H is inhibited by non-phosphorylated reaction products. Alphafold (11) modeling helped us identify a potential allosteric Rab GTPase binding site located behind the PPM1H active site. We present here our biochemical characterization of end-product, allosteric inhibition of PPM1H phosphatase.

## Results and Discussion

Figure 1 shows an Alphafold model (11) for PPM1H bound to phosphoRab10 in the presence of non-phosphorylated Rab10 protein. The structure of PPM1H was first reported by Amir Khan and colleagues (12); Alphafold correctly orients the phosphorylated Rab10 such that the substrate phosphate points directly into the enzyme active site. [Note that PPM1H is a dimer (12); we show the interactions for one half of the dimeric model for clarity. None of these interactions would be obscured in a dimer model.] Significant surface interaction is also seen between the FLAP domain and the phosphoRab substrate, consistent with the role of the FLAP in substrate selectivity. In this representation we have deleted loops #1 and #2 (residues 115-133 and 204-217) that do not influence protein localization or enzyme activity (10) and were not resolved in the crystal structure ((12) see Fig. S1A). The PPM1H Leu66 sidechain is predicted to contribute to non-phosphorylated Rab10 binding. Also note the location of the PPM1H N-terminal alpha helix that will be discussed below.

**Figure 1:**
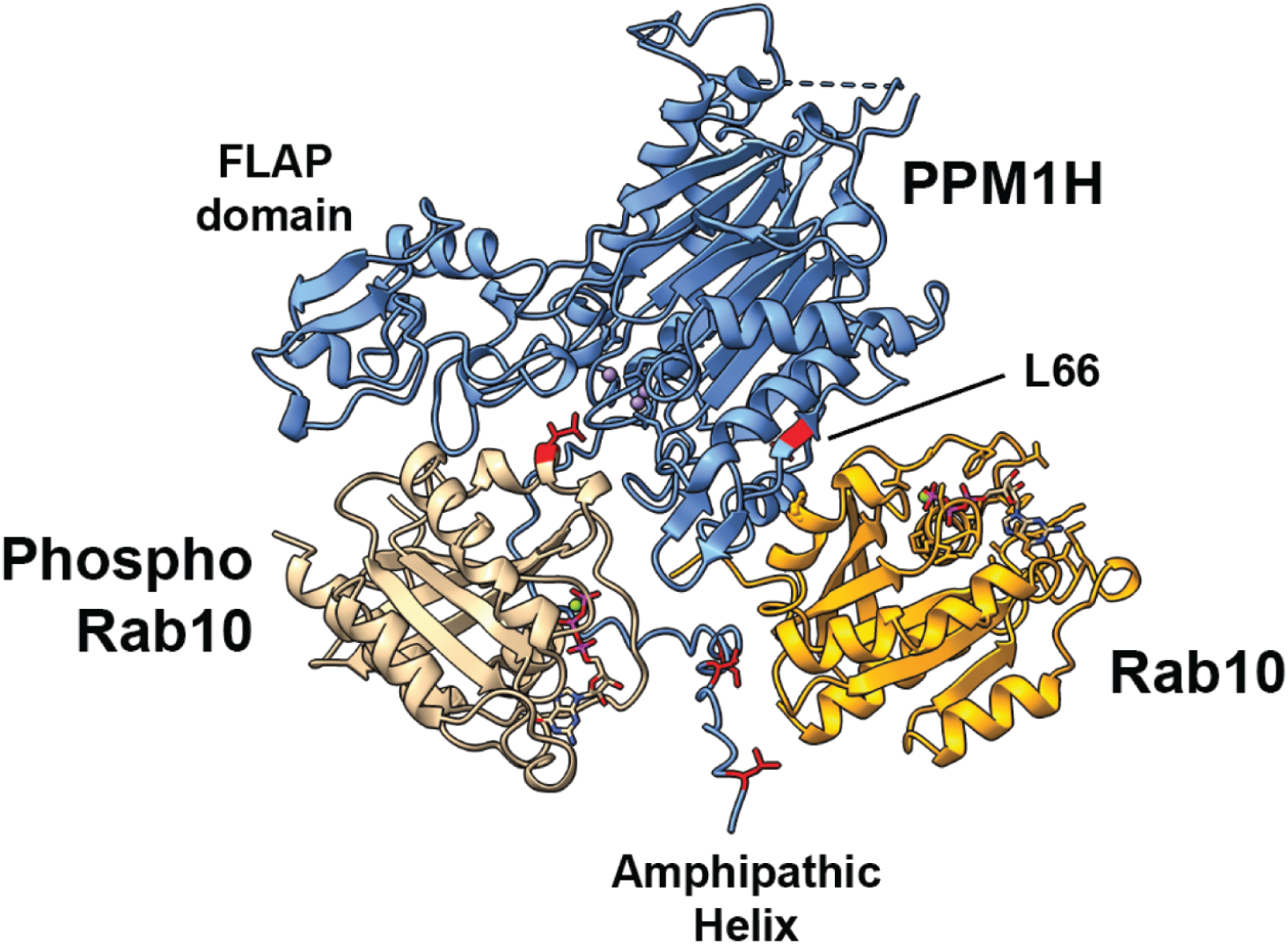
Alphafold (11) model of PPM1H bound to its phosphoRab10 substrate and non-phosphoRab10 allosteric regulator (loops #1 and #2 were removed for simplicity (dashed lines); see Figure S1A for comparison), created using Chimera software (13). PPM1H (blue); phosphoRab10 (beige); Rab10 (goldenrod). The PPM1H FLAP domain (12), amphipathic helix (10) and Leu66 residue are indicated. Also colored are amphipathic helix residues shown to be important for liposome binding and localization of PPM1H in cells (10).

To test the validity of the Alphafold model, we engineered a fluorescent version of full length PPM1H in which we inserted mNeon into loop#1 at position 125. Alphafold modeling of PPM1H with and without this insertion (Fig. S1) indicated that the addition of mNeon would not alter overall PPM1H protein structure. HisSUMO-mNeon PPM1H, HisSUMO-mNeon PPM1H L66R and HisSUMO-mNeon Δ37 PPM1H were expressed in bacteria and purified by Cobalt affinity and size exclusion chromatography for further analysis; the HisSUMO portions were removed by Ulp1 SUMO protease cleavage prior to gel filtration (Fig. S1B-D).

Figure 2 compares the binding of PPM1H to Rab10 (Fig. 2A), Rab8A (Fig. 2B), and Rab12 (Fig. 2C) monitored using microscale thermophoresis (MST). Non-phosphorylated Rab10 and Rab8A bound PPM1H with a KD of 1.3 and 1.4 µM respectively, while Rab12 bound much more weakly with a KD of 7.4 µM (Table 1) which would not be detected in cells. When the same Rabs were tested for binding to PPM1H L66R, the KD values increased about 3 fold to 4.4 µM and 5.2 µM for Rabs10 and 8A (Fig. 2D,E; Table 1); binding of Rab12 weakened further to 9.6 µM (Fig. 2F; Table 1). Thus, the L66R mutation blocks non-phosphoRab binding to PPM1H.

**Table 1:** Summary of PPM1H binding affinities (KD)

|  | mNeon PPM1H (μM) | mNeon L66R PPM1H (μM) | mNeon Δ37 PPM1H (μM) |
| --- | --- | --- | --- |
| Rab8A | 1.4 ± 0.98 | 5.2 ± 3 | 10.5 ± 6 |
| Rab10 | 1.3 ± 0.9 | 4.4 ± 2.7 | 6.5 ± 4.5 |
| Rab12 | 7.4 ± 3.8 | 9.6 ± 5.1 |  |
| thio-pRab8A | 0.99 ± 0.63 | 1.1 ± 0.76 |  |

**Figure 2:**
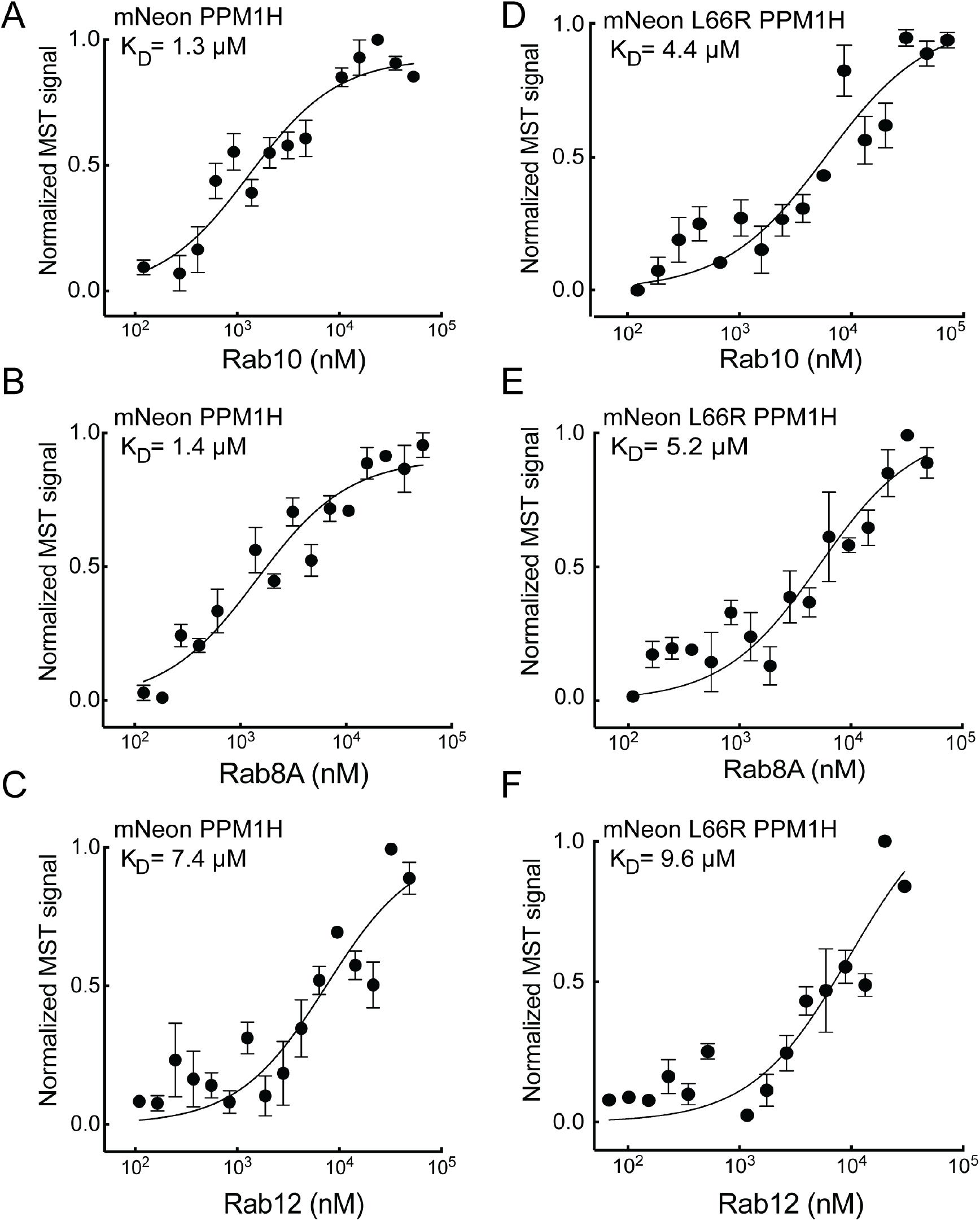
PPM1H binds non-phosphorylated Rab8A and Rab10 but not Rab12 via Leu66. Microscale thermophoresis of mNeon PPM1H and mNeon L66R PPM1H with Rab10 (A, B), Rab8A (C, D), or Rab12 (E, F). Purified Rab proteins were serially diluted, and mNeon PPM1H or mNeon L66R PPM1H was added (final concentration 100 nM). Graphs show the mean and SEM from 3 independent measurements, each using different protein preparations. Values are summarized in Table 1.

To ensure that the non-phosphoRab binding site is distinct from that used by phosphoRab substrates, we used a poorly hydrolyzable, ATP*γ*S thio-phosphorylated Rab substrate to enable estimation of the affinity of PPM1H for its phosho-Rab substrate. PhosTag gels showed that the protein was fully modified for use in this experiment and resisted PPM1H action in a 15 minute in vitro reaction (Fig. S2).

Thiophosphorylated Rab8A bound PPM1H with an affinity of 0.99µM (Fig. 3A); importantly, the KD was not altered when tested with PPM1H L66R protein (KD = 1.1 µM; Fig. 3B). These experiments confirm that PPM1H binds non-phosphorylated Rabs via at least one residue (L66) that is not needed for phosphoRab substrate binding, consistent with the Alphafold model (Fig. 1).

**Figure 3:**
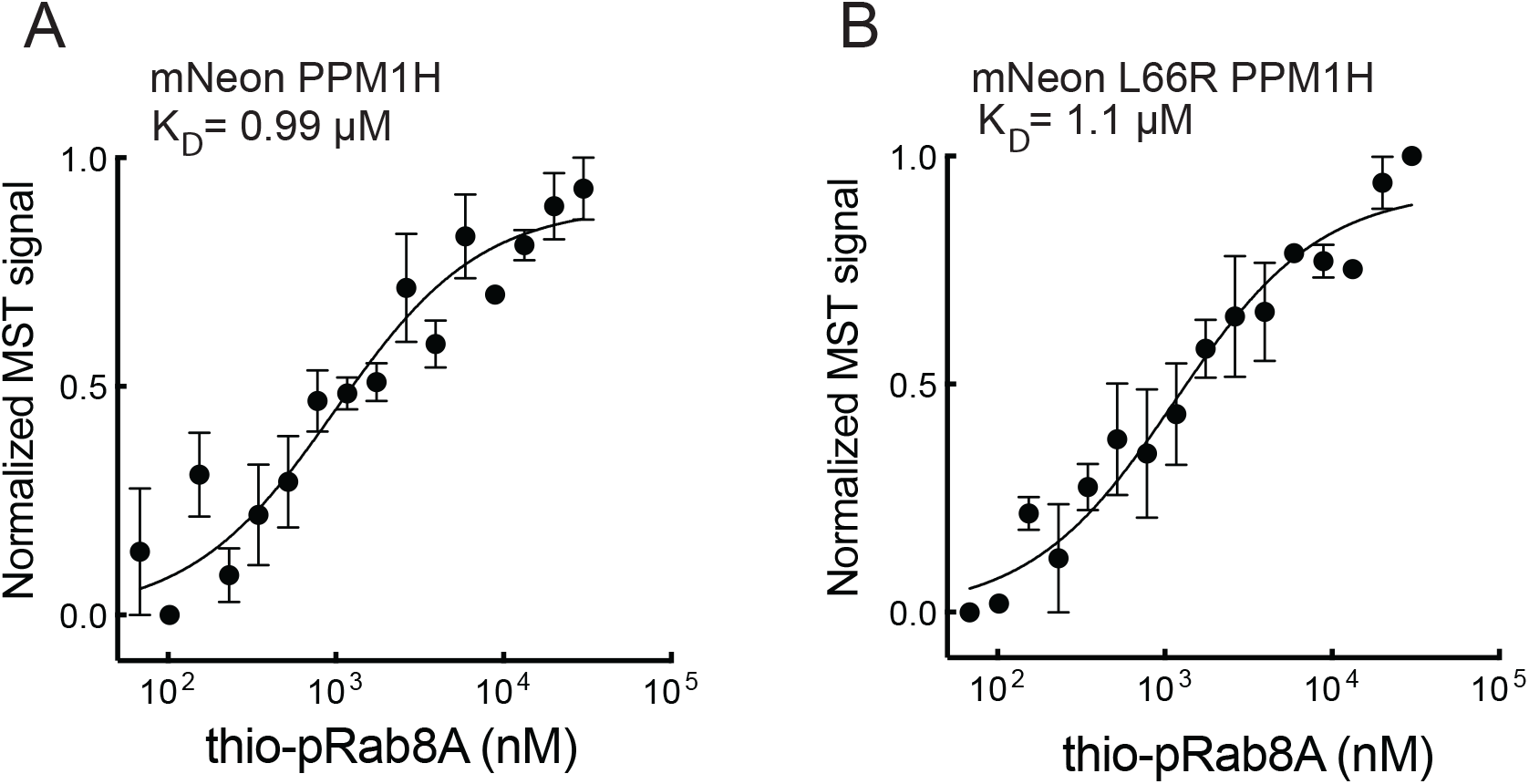
PPM1H L66 mutation does not disrupt phosphoRab substrate binding. Microscale thermophoresis of mNeon PPM1H (A) or mNeon PPM1H L66R (B) bound to thio-phosphorylated Rab8A as in Fig. 2.

In previous work we showed that full length PPM1H protein floats to the top of a sucrose gradient in the presence of liposomes prepared using a Golgi-type lipid composition (10). As an independent test of protein:protein interaction, we utilized this method to ask whether liposome-bound PPM1H would be sufficient to cause a non-prenylated, non-phosphorylated Rab binding partner to also float to the top of a sucrose gradient. Figure 4 shows examples of this type of experiment. mNeon PPM1H remained at the bottom of the gradient unless it was allowed to bind to liposomes; similarly, non-prenylated Rab8A was detected at the bottom of the tube unless both PPM1H and liposomes were added (Fig. 4A,B). Importantly, Rab8A failed to float with liposomes bearing PPM1H L66R (Fig. 4C,D), despite the ability of that PPM1H mutant to bind and float with liposomes. These experiments confirm that non-phosphorylated Rab8A interacts with PPM1H in a manner that requires PPM1H L66.

**Figure 4:**
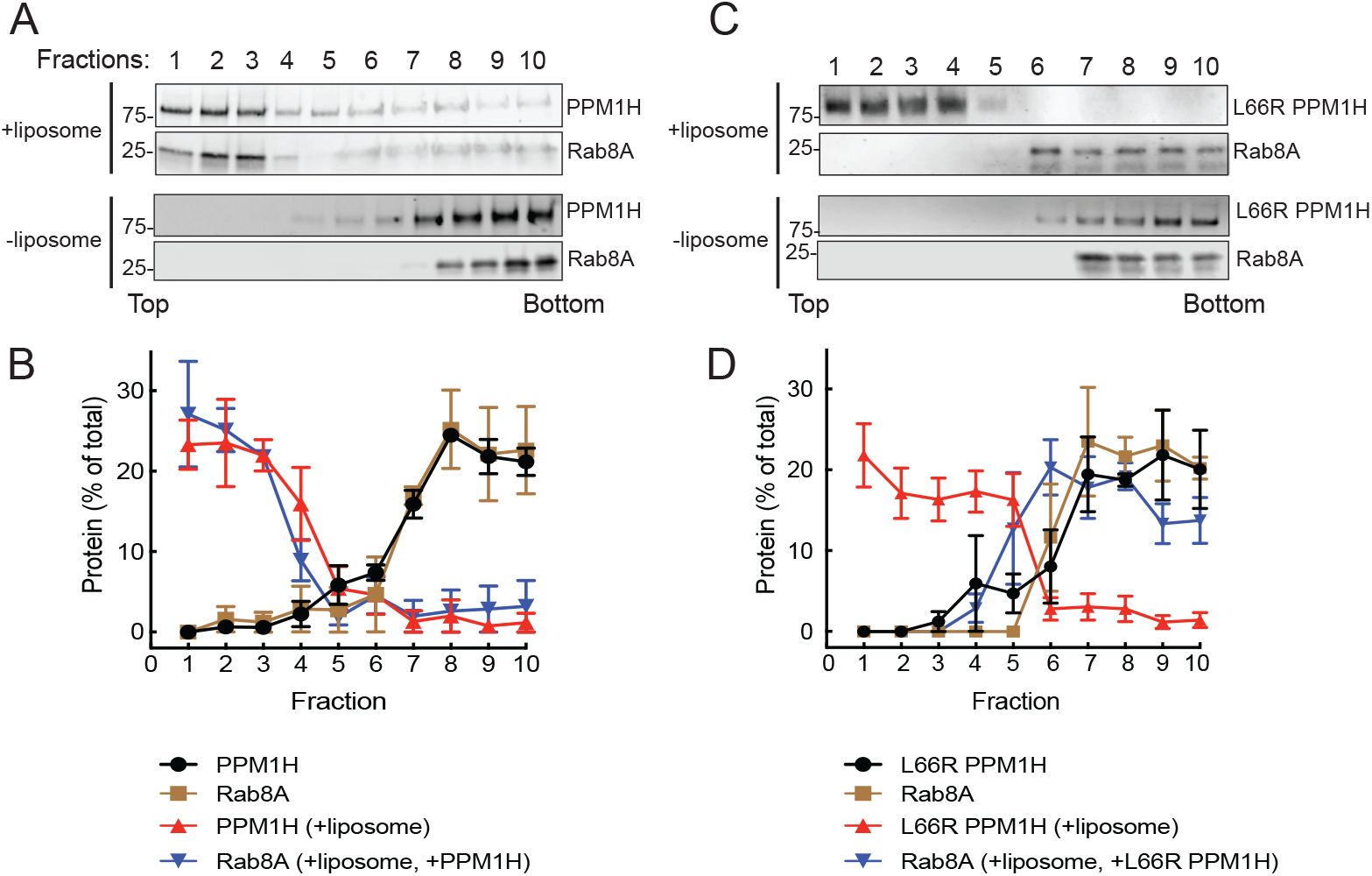
Rab8A associates with liposome-bound PPM1H. Sucrose gradient co-flotation of His-Rab8A (full length, non-prenylated) with mNeon PPM1H (A) or mNeon L66R PPM1H (B) in the presence and absence of 50nm liposomes. The distribution of PPM1H and Rab8A across the gradient (B,D) was determined by immunoblot (A,C); fractions were collected from the top. Quantification of three independent experiments is shown (±SEM). Numbers at the edge of each blot in this and all subsequent figures indicate molecular weight marker mobility shown in kDa.

Identical results were obtained in experiments analyzing the ability of PPM1H to bind to non-phosphorylated Rab10 (Fig. 5A-D). In contrast, non-phosphorylated Rab12 failed to float with liposome-bound wild type or mutant PPM1H (FIg. 6A-D), consistent with the low KD determined by MST (Fig. 2C). Thus, Rab8A and Rab10, but not Rab12, can bind to PPM1H is a manner that requires PPM1H L66.

**Figure 5:**
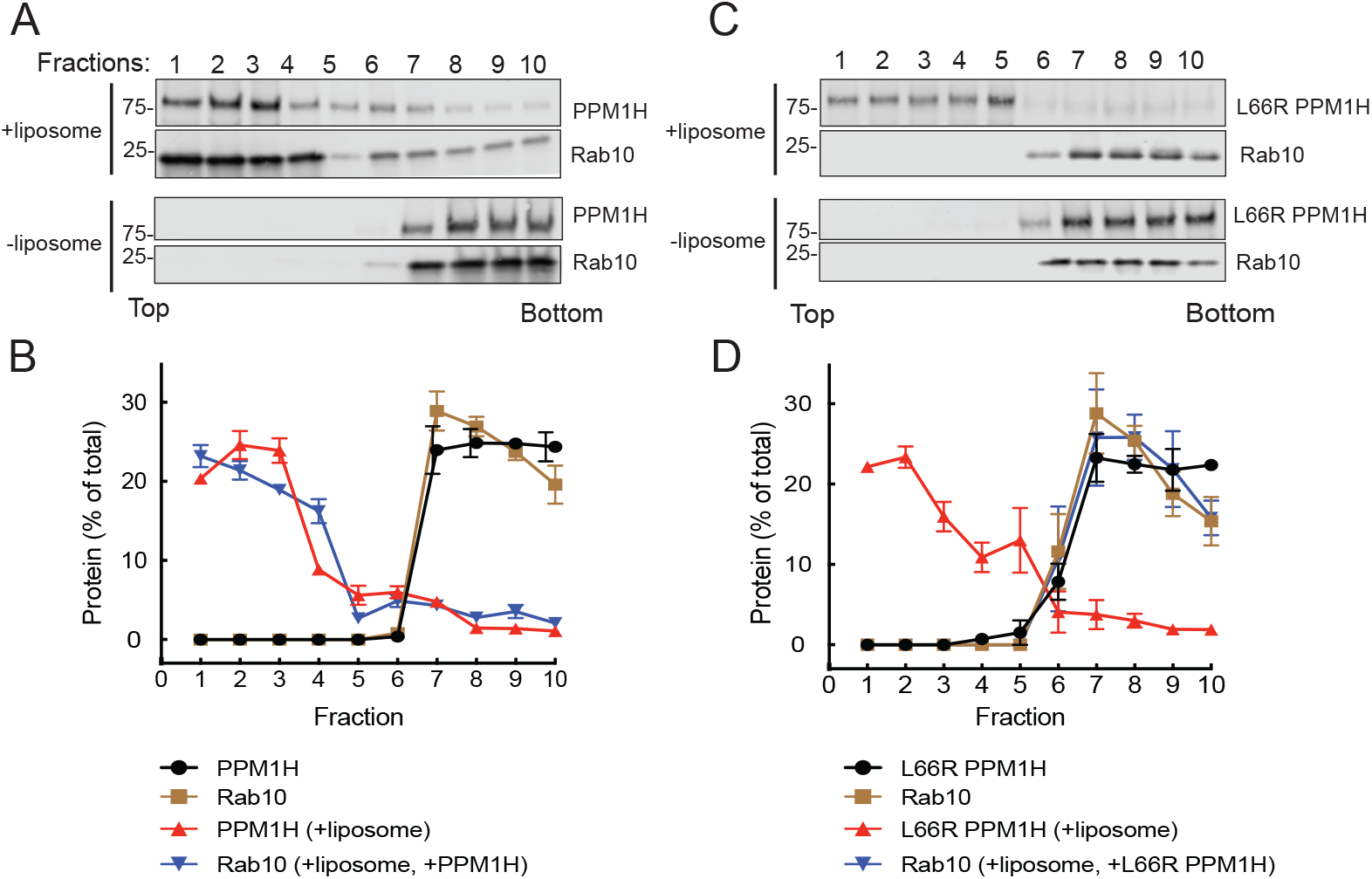
Rab10 associates with liposome-bound PPM1H. Sucrose gradient co-flotation of Rab10 (full length, non-prenylated) with mNeon PPM1H (A) or mNeon L66R PPM1H (B) in the presence and absence of 50nm liposomes. The distribution of PPM1H and Rab10 across the gradient (B,D) was determined by immunoblot (A,C); fractions were collected from the top. Quantification of three independent experiments is shown (±SEM).

**Figure 6:**
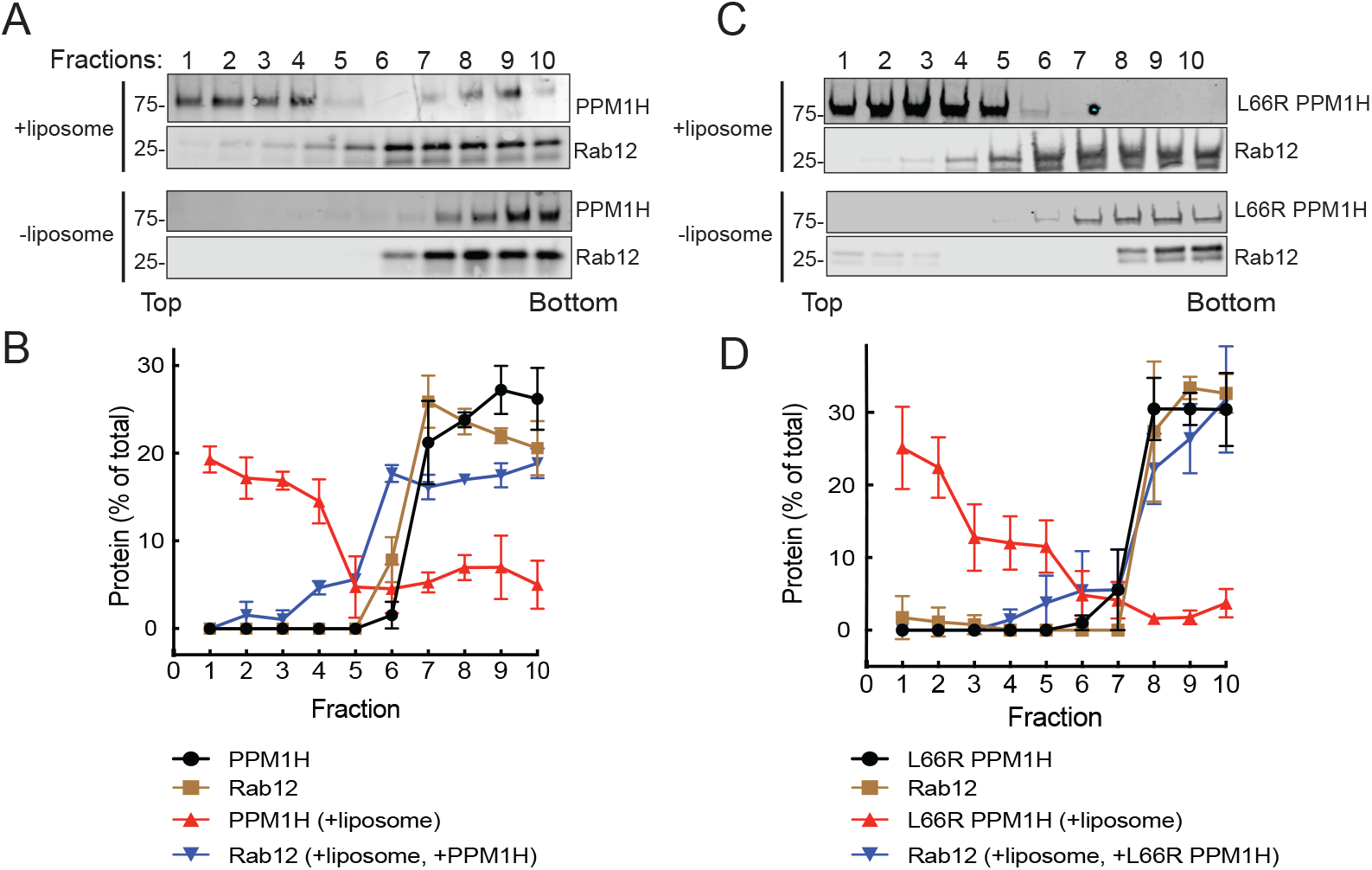
Rab12 fails to associate with liposome-bound PPM1H. Sucrose gradient co-flotation of Rab12 (full length, non-prenylated) with mNeon PPM1H (A) or mNeon L66R PPM1H (B) in the presence and absence of 50nm liposomes. The distribution of PPM1H and Rab12 across the gradient (B,D) was determined by immunoblot (A,C); fractions were collected from the top. Quantification of three independent experiments is shown (±SEM).

### End-product inhibition of PPM1H

Rab10 dephosphorylation was monitored in vitro using purified proteins in conjunction with immunoblot analysis using a phosphoRab10-specific antibody. As shown in Figure 7A, phosphoRab10 was readily detected and as expected, its abundance decreased upon addition of purified PPM1H enzyme. Fig. 7 panels B and C show re-plots of the data to display PPM1H *activity* rather than loss of pRab10 protein: full activity is defined as the amount of pRab remaining in the presence of 100nM PPM1H protein (lanes 3 and 4, Fig. 7A). When reactions also contained micromolar amounts of unphosphorylated Rab8A protein, a concentration-dependent inhibition of phosphatase activity was observed (Fig. 7A,B). In contrast, addition of comparable amounts of Rab12 failed to inhibit PPM1H enzyme (Fig. 7C), consistent with the very weak interaction of Rab12 with PPM1H (Figs. 2, 6). The concentrations of Rab8A or Rab10 needed to achieve 50% inhibition were entirely consistent with KD determinations (Figure 2; Table 1). Thus, PPM1H displays end-product inhibition by its cellular Rab8A and Rab10 substrates. Note that Rab12 is not an endogenous, cellular substrate for PPM1H enzyme (6, 7).

**Figure 7:**
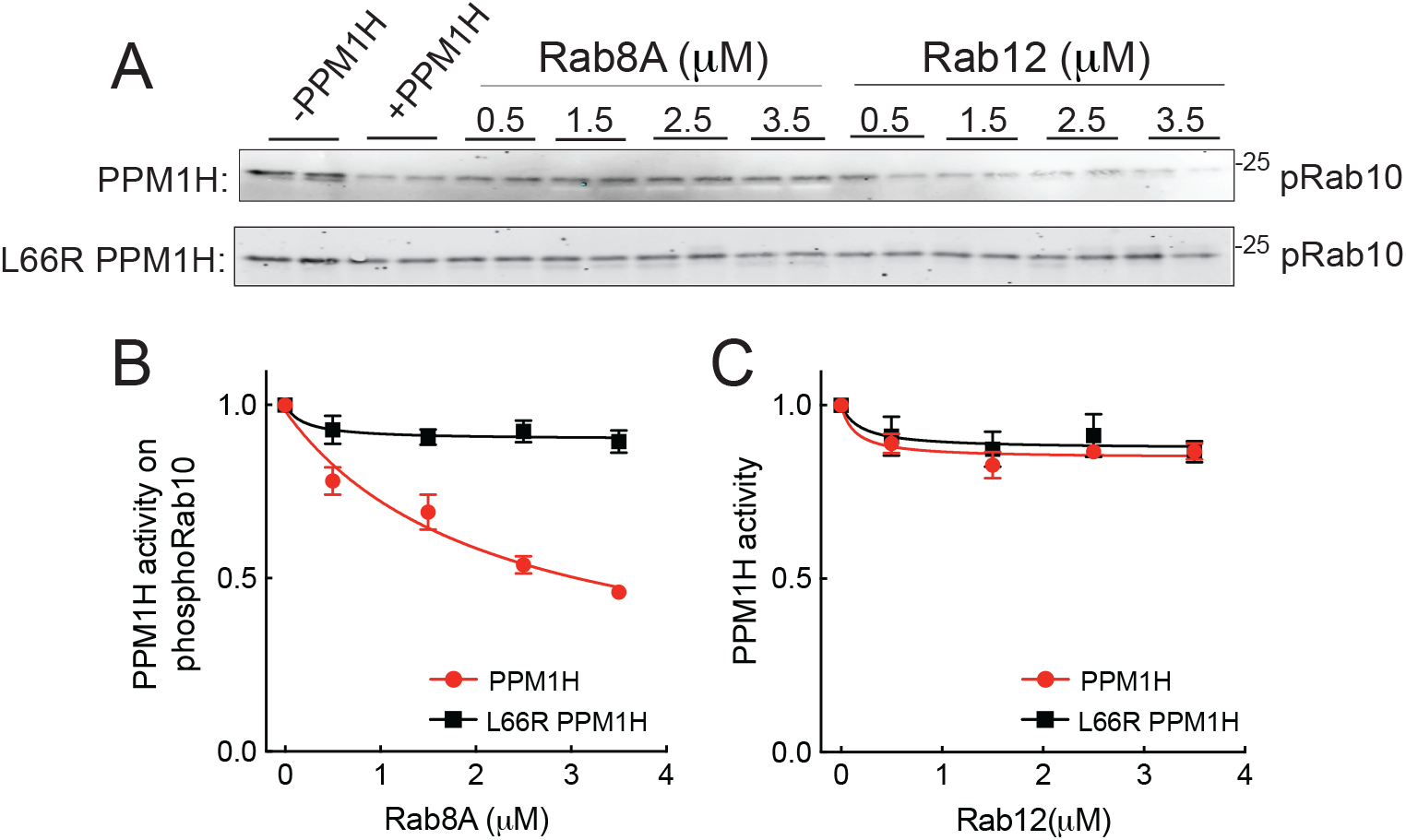
Rab8A but not Rab12 inhibits PPM1H. Purified mNeon PPM1H or mNeon L66R PPM1H was assayed for phosphoRab10 phosphatase activity in the presence of Rab8A full length (left) or Rab12 full length (right) for 15 minutes plus 50 nm liposomes to activate the enzyme. A. Anti-pRab10 immunoblot for reactions with PPM1H (upper gel) or L66R PPM1H (lower gel). B,C. Data were plotted as PPM1H activity, with full activity equal to the amount of dephosphorylation seen in lanes 3 and 4 of each gel, compared with reactions lacking PPM1H (lanes 1 and 2). Error bars represent SEM from three independent experiments; representative gels are shown. Black line, mNeon L66R PPM1H activity; red line, mNeon PPM1H activity.

### Non-phosphorylated Rabs require PPM1H N-terminal sequences for binding

We showed previously that deletion of the N-terminal 37 residues caused PPM1H to display an entirely cytosolic localization (10). Thus, at first, it was hard to explain why our newly discovered, relatively high affinity Rab binding site was not sufficient to localize PPM1H to Rab8A and Rab10-positive Golgi membranes in cells. One possible explanation was that the PPM1H N-terminus also contributes to Rab8A and Rab10 binding. To test this, we compared the binding of non-phosphorylated Rab proteins to wild type and Δ37-PPM1H. To our surprise, binding of Rab8A and Rab10 to Δ37-PPM1H missing the amphipathic helix decreased from 1.3 and 1.4µM, respectively to 10.5 and 6.5µM (Fig. 8A,B; Table 1). Thus, in addition to L66, PPM1H’s N-terminal 37 residues are also needed for both liposome association and non-phosphoRab protein binding.

**Figure 8:**
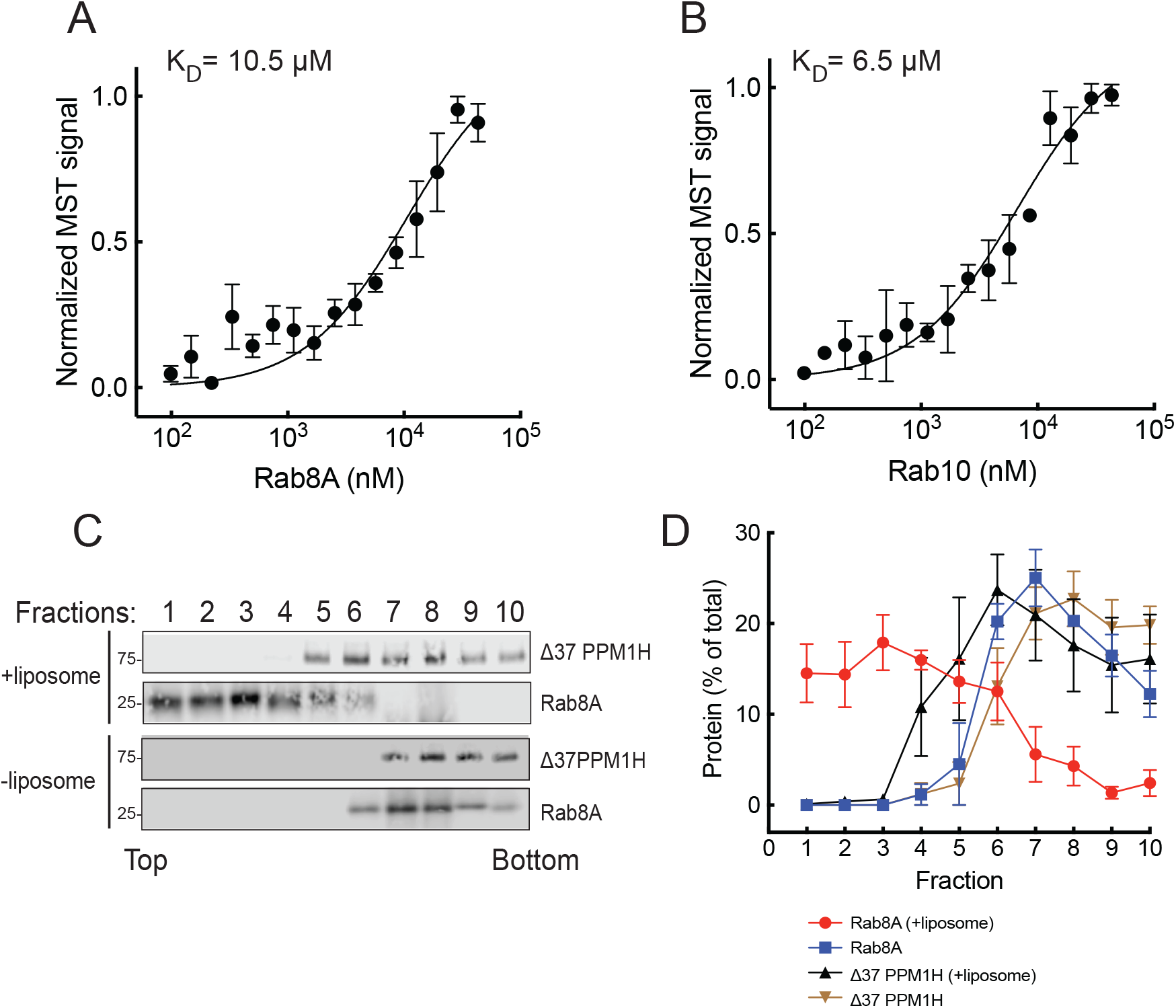
PPM1H’s N-terminal 37 residues contribute to Rab8A and Rab10 binding. Microscale thermophoresis of mNeon Δ37 PPM1H with Rab8A (A) or Rab10 (B). C, D. Sucrose gradient co-flotation of Rab8A with mNeon Δ37 PPM1H in the presence and absence of 50nm DGS NTA(Ni) (Nickel-Nitrilotriacetic acid) containing liposomes as indicated, as in Fig. 4.

As an independent measure of the importance of the PPM1H N-terminus in Rab binding, we artificially anchored Rab8A-6XHis onto the surface of NTA-lipid containing liposomes and tested whether binding at the PPM1H allosteric site would be sufficient to float Δ37 PPM1H. As shown in Fig. 7C, despite efficient flotation of Rab8A-6XHis with the NTA-lipid-containing liposomes, Δ37 PPM1H failed to float to the top of the sucrose gradient, consistent with its weak affinity for Rab8A determined by MST. Altogether these experiments confirm the importance of the PPM1H N-terminus, both for binding to highly curved membranes and to non-phosphorylated Rab proteins. It is not alone sufficient for binding because L66R PPM1H also failed to bind Rab8A and Rab10 proteins, despite the presence of the PPM1H N-terminus. Because PPM1H is a dimer, these results might be explained if one N-terminal sequence engages the liposome while the other engages the Rab GTPase; alternatively, it may be possible that one face of the amphipathic helix can be anchored in the liposome membrane while its exposed surface interacts with Rab proteins.

In summary, we have reported here the existence of an allosteric binding site for non-phosphorylated, substrate Rab GTPases on PPM1H enzyme and shown that binding at this site inhibits PPM1H catalytic activity. In addition to regulating PPM1H activity, Rab binding appears to contribute to membrane localization of PPM1H via the phosphatase’s N-terminal amphipathic helix, since Rab binding and localization both rely on this sequence. Does this mean that membrane-associated PPM1H is held in an inactive conformation? We showed previously that in the absence of non-phosphorylated Rab proteins, PPM1H is activated upon amphipathic helix binding to highly curved (50nm) liposomes (10). In that study, we concluded that such binding would enrich the enzyme at highly curved Golgi rims. We speculate that an initial Rab protein interaction brings PPM1H to the correct membrane for substrate de-phosphorylation; perhaps an equilibrium between Rab-bound versus highly curved membrane bilayer-bound forms of the enzyme regulates PPM1H activity in a spatially controlled manner. Altogether, these findings identify a dual role for Golgi-localized, non-phosphorylated Rab GTPases in the spatial localization and regulation of PPM1H, revealing a layer of phosphatase control that could be leveraged to enhance dephosphorylation of LRRK2 substrates.

## Materials and Methods

### Cloning and plasmids

DNA constructs were amplified in *Escherichia coli* DH5α and purified using mini prep columns (EconoSpin). DNA sequencing was performed by MC LAB (https://mclab.com/). pET15b His-MST3 was a generous gift from Amir Khan (Trinity College, Dublin). mNeon was cloned into loop1 (residues 115-133) of pET15b His SUMO PPM1H and pET15b His SUMO Δ37 PPM1H backbones. The PPM1H L66R point mutation was introduced using site-directed mutagenesis. His Rab8A Q67L was subcloned from HA-Rab8A (DU35414, MRC PPU) into the pET14b vector. His SUMO Rab10 (full length) Q68L and His SUMO Rab12 (full length) Q101L were cloned from pET15b Rab10 Q68L and pQE 80L Rab12 (Q101L), respectively, into the pET15b backbone. All cloning and subcloning was performed using Gibson assembly (https://www.protocols.io/view/gibson-assembly-cloning-eq2lyjwyqlx9/v1).

### Protein Purification

The following protocol describes purification of His SUMO mNeon PPM1H, His SUMO mNeon Δ37 PPM1H, His SUMO mNeon L66R PPM1H, His SUMO Rab10 (Q68L), His Rab8A (Q67L), His MST3, and His SUMO Rab12 (Q101L) https://www.protocols.io/view/ppm1h-purification-from-e-coli-d67v9hn6. All Rab proteins were full length and non-prenylated. Rab Q mutants were used throughout this study; these favor the GTP-bound state, making it easier to ensure that the Rabs are as active as possible in these experiments. For Rab protein purification, 20µM GTP was included in all buffers. Briefly, bacterial cultures were grown at 37°C in Luria Broth and induced with 0.3 mM isopropyl-1-thio-β-d-galactopyranoside when the optical density at 600 nm reached 0.5–0.6. Cultures were harvested after overnight growth at 18°C. The cell pellets were resuspended in ice-cold lysis buffer containing 50 mM HEPES (pH 8.0), 10% glycerol, 500 mM NaCl, 10 mM imidazole, 5 mM MgCl2, 0.2 mM TCEP, 20 μM GTP, and an EDTA-free protease inhibitor cocktail (Roche #69040300). Bacterial cells were lysed using an Emulsiflex-C5 apparatus (Avestin) at 10,000 psi and centrifuged at 40,000 rpm for 45 minutes at 4°C using a Beckman Ti45 rotor. The clarified lysate was filtered through a 0.2 µm filter (Nalgene) and passed over a HiTrap TALON crude 1 mL column (Cytiva #28953766). The column was washed with lysis buffer until the absorbance returned to baseline, and proteins were eluted using a gradient of 100–500 mM imidazole in the same buffer. Peak fractions were identified by SDS-PAGE using 4–20% Precast Protein Gels. For His-MST3 (Mammalian STE20-like protein kinase 3) and His Rab8A, the eluted proteins were concentrated by Centricon filtration before being applied to a Superdex™ 75 10/300 GL column (Cytiva #29148721).

TALON purified, His-SUMO-tagged proteins were dialyzed overnight together with Ulp1 SUMO protease (100 ng/mg of protein) to remove the His-SUMO tag. The following day, the dialyzed proteins were collected, concentrated, and applied onto a Superdex™ 200 10/300 GL column (Cytiva #28990944) fitted in series with an additional 1 mL HiTrap TALON crude column (Cytiva #28953766) to remove uncleaved His-SUMO protein, the His-tagged Ulp1 SUMO protease, and the HisSUMO tag.

### Microscale Thermophoresis

Protein–protein interactions were analyzed using Microscale Thermophoresis (MST) on a Monolith NT.115 instrument (NanoTemper Technologies) as described in detail: <u>dx.doi.org/10.17504/protocols.io.bvvmn646</u>. His-tagged Rab8A Q67L was incubated overnight at 4°C with His-tagged MST3 kinase in a reaction buffer containing 50 mM HEPES (pH 8), 100 mM NaCl, 5 mM MgCl2, 2 mM ATP or ATPγS, 100 μM GTP, 0.5 mM TCEP, 10% glycerol, and 5 μM BSA. In all experiments, the unlabeled (Rab12 Q101L, Rab10 Q68L, His Rab8A Q67L, His pRab8A Q67L) protein partner was titrated against a fixed concentration of fluorescent mNeon PPM1H, mNeon Δ37 PPM1H or mNeon L66R PPM1H (100 nM). Sixteen serial dilutions of the unlabeled protein partner were prepared to generate a complete binding isotherm. Binding reactions were carried out in a reaction buffer in 0.5 mL Protein LoBind tubes (Eppendorf) and incubated in the dark for 30 minutes before being loaded into NT.115 premium treated capillaries (NanoTemper Technologies).

Measurements were taken using a blue LED set at 30% excitation power (excitation: 450–480 nm; emission: 515–570 nm) and an IR-laser at 60% power for 30 seconds, followed by 5 second of cooling. Data were analyzed using NTAffinityAnalysis software (NanoTemper Technologies), where raw fluorescence data were used to generate binding isotherms. These isotherms were fitted using both NanoTemper software and GraphPad Prism to calculate the dissociation constant (KD) via nonlinear regression. Binding affinities derived from both methods were consistent.

### Liposome Preparation

A detailed step-by-step protocol can be found at: https://doi.org/10.17504/protocols.io.5qpvo3ke7v4o/v1. Briefly, liposomes were prepared using a Golgi-like lipid composition consisting of (18:0-20:4)PC, (18:0-20:4)PI, (18:0-18:2)PS, (18:1) plus PI(4)P, and cholesterol in a molar ratio of 78:7:5:1:9 (all lipids from Avanti Polar Lipids). For the flotation assay of His-Rab8A (Q67L) with mNeon Δ37 PPM1H, the liposome mixture included (18:1)DGS NTA (Ni2+) and was prepared using a molar ratio of 76:7:5:1:9:2.

The lipid mixtures, dissolved in chloroform, were dried under a nitrogen stream to form a thin film. This film was then resuspended in 50 mM HEPES (pH 7.5) and 120 mM KCl. The suspension underwent two brief 5-second sonication cycles in a bath sonicator and was subsequently extruded through 0.4 μm, 0.1 μm, and 0.05 μm polycarbonate filters (21 passes each) using a hand extruder (Avanti). The final liposome suspension had a lipid concentration of 15 mM.

### Sucrose Density Gradient Co-flotation

Liposome co-flotation assays were conducted by incubating 3µM Rab10, Rab12, or His Rab8A with 1.5 mM, 50nm liposomes and 1µM mNeon PPM1H, mNeon L66R PPM1H or mNeon Δ37 PPM1H for 30 minutes at 30°C in a total volume of 33 µL using HKM buffer (20 mM HEPES, pH 7.5, 150 mM potassium acetate, 1 mM magnesium chloride, 20 µM GTP). Next, 167 µL of 60% (w/v) sucrose was added and mixed to adjust the solution to 50% sucrose. The sample was then overlaid with 200 µL of 25% (w/v) sucrose and 100 µL of HKM buffer. The samples were centrifuged at 45,000 RPM for 3h using a Ti55 swinging bucket rotor. Ten fractions (50 µL each) were manually collected from the top of the gradient using a micropipette and analyzed by immunoblotting using a rabbit anti-PPM1H antibody. Rab10, Rab12, and Rab8A were detected using mouse anti-Rab10, anti-Rab12, and anti-Rab8A antibodies, respectively.

### In vitro PPM1H phosphatase assay

A detailed protocol is available: <u>dx.doi.org/10.17504/protocols.io.5jyl8d4j7g2w/v1</u>. Briefly, untagged Rab10 Q68L (full-length) was incubated overnight at 4°C with His-MST3 kinase in a reaction buffer containing 50 mM HEPES (pH 8), 100 mM NaCl, 5 mM MgCl2, 2 mM ATP, 100 μM GTP, 0.5 mM TCEP, 10% glycerol, and 5 μM BSA to achieve Rab10 phosphorylation. The following day, His-MST3 kinase was removed by passing the reaction mixture through a 1mL syringe column containing 100 μL (50%) Ni-NTA slurry, and the flowthrough containing phosphorylated Rab10 was collected. Next, 1.5 μM phosphorylated Rab10 was incubated with 100 nM mNeon PPM1H (or 100 nM mNeon L66R PPM1H), in the presence of 0.5 μM, 1.5 μM, 2.5 μM, and 3.5 μM of unphosphorylated Rab8A and Rab12 with 50nM liposomes, at 30°C for 15 minutes. The reactions were stopped by adding a 5X SDS-PAGE sample buffer. Samples were analyzed by immunoblotting to assess Rab10 dephosphorylation using anti-pRab10 (1:1,000) antibody.

### Data Analysis

Data analysis was carried out using GraphPad Prism version 9 for Mac, GraphPad Software, Boston, Massachusetts USA, www.graphpad.com.

## Abbreviations

LRRK2: Leucine rich repeat kinase
MST: microscale thermophoresis
MST3: Mammalian STE20-like protein kinase 3
NTA(Ni): Nickel-Nitrilotriacetic acid

## Data and materials availability

All reagents used in this study are available from commercial sources or repositories without restriction and RRIDs for all reagents are provided in the Key Resource Table 2. All primary data have been deposited in Zenodo and can be found at https://doi.org/10.5281/zenodo.15427865.

**Table 2:**
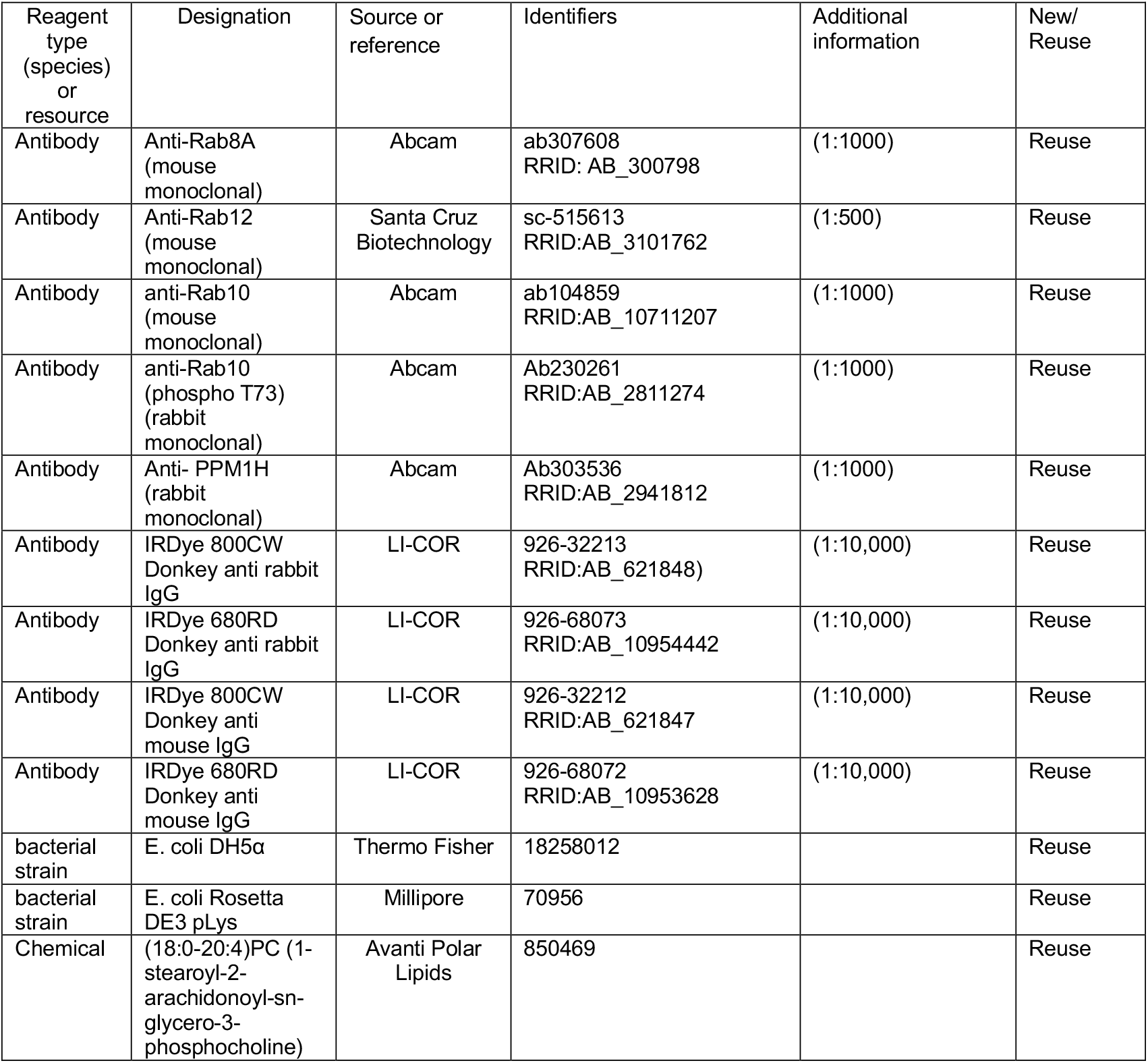
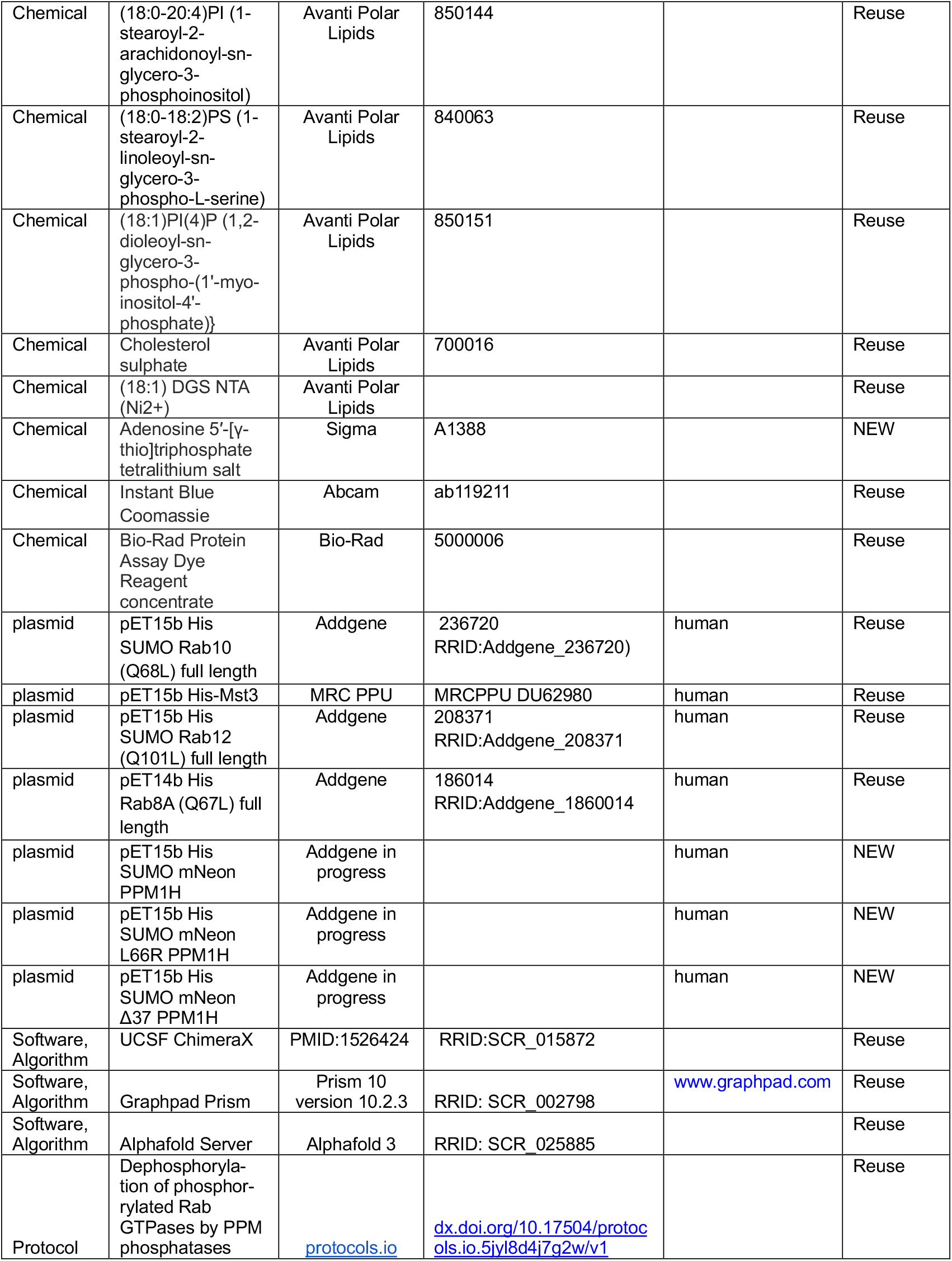
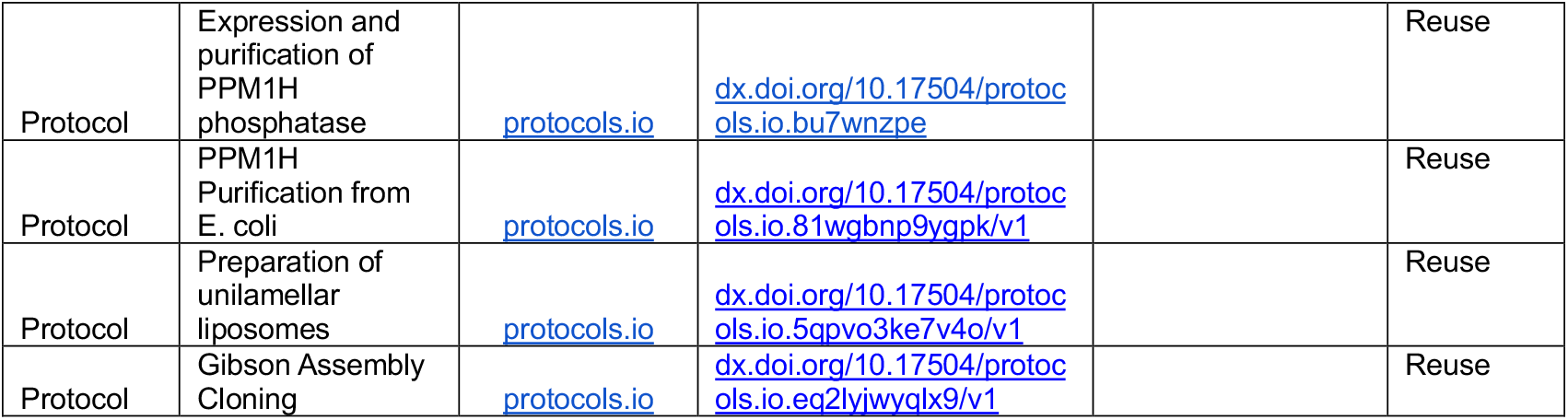
Key Resources

| Reagent type (species) or resource | Designation | Source or reference | Identifiers | Additional information | New/ Reuse |
| --- | --- | --- | --- | --- | --- |
| Antibody | Anti-Rab8A (mouse monoclonal) | Abcam | ab307608<br>RRID: AB_300798 | (1:1000) | Reuse |
| Antibody | Anti-Rab12 (mouse monoclonal) | Santa Cruz Biotechnology | sc-515613<br>RRID:AB_3101762 | (1:500) | Reuse |
| Antibody | anti-Rab10 (mouse monoclonal) | Abcam | ab104859<br>RRID:AB_10711207 | (1:1000) | Reuse |
| Antibody | anti-Rab10 (phospho T73) (rabbit monoclonal) | Abcam | Ab230261<br>RRID:AB_2811274 | (1:1000) | Reuse |
| Antibody | Anti- PPM1H (rabbit monoclonal) | Abcam | Ab303536<br>RRID:AB_2941812 | (1:1000) | Reuse |
| Antibody | IRDye 800CW Donkey anti rabbit IgG | LI-COR | 926-32213<br>RRID:AB_621848) | (1:10,000) | Reuse |
| Antibody | IRDye 680RD Donkey anti rabbit IgG | LI-COR | 926-68073<br>RRID:AB_10954442 | (1:10,000) | Reuse |
| Antibody | IRDye 800CW Donkey anti mouse IgG | LI-COR | 926-32212<br>RRID:AB_621847 | (1:10,000) | Reuse |
| Antibody | IRDye 680RD Donkey anti mouse IgG | LI-COR | 926-68072<br>RRID:AB_10953628 | (1:10,000) | Reuse |
| bacterial strain | E. coli DH5α | Thermo Fisher | 18258012 |  | Reuse |
| bacterial strain | E. coli Rosetta DE3 pLys | Millipore | 70956 |  | Reuse |
| Chemical | (18:0-20:4)PC (1-stearoyl-2-arachidonoyl-sn-glycero-3-phosphocholine) | Avanti Polar Lipids | 850469 |  | Reuse |
| Chemical | (18:0-20:4)PI (1-stearoyl-2-arachidonoyl-sn-glycero-3-phosphoinositol) | Avanti Polar Lipids | 850144 |  | Reuse |
| Chemical | (18:0-18:2)PS (1-stearoyl-2-linoleoyl-sn-glycero-3-phospho-L-serine) | Avanti Polar Lipids | 840063 |  | Reuse |
| Chemical | (18:1)PI(4)P (1,2-dioleoyl-sn-glycero-3-phospho-(1'-myo-inositol-4'-phosphate)) | Avanti Polar Lipids | 850151 |  | Reuse |
| Chemical | Cholesterol sulphate | Avanti Polar Lipids | 700016 |  | Reuse |
| Chemical | (18:1) DGS NTA (Ni<sup>2+</sup>) | Avanti Polar Lipids |  |  | Reuse |
| Chemical | Adenosine 5'-[γ-thio]triphosphate tetralithium salt | Sigma | A1388 |  | NEW |
| Chemical | Instant Blue Coomassie | Abcam | ab119211 |  | Reuse |
| Chemical | Bio-Rad Protein Assay Dye Reagent concentrate | Bio-Rad | 5000006 |  | Reuse |
| plasmid | pET15b His SUMO Rab10 (Q68L) full length | Addgene | 236720<br>RRID:Addgene_236720) | human | Reuse |
| plasmid | pET15b His-Mst3 | MRC PPU | MRCPPU DU62980 | human | Reuse |
| plasmid | pET15b His SUMO Rab12 (Q101L) full length | Addgene | 208371<br>RRID:Addgene_208371 | human | Reuse |
| plasmid | pET14b His Rab8A (Q67L) full length | Addgene | 186014<br>RRID:Addgene_1860014 | human | Reuse |
| plasmid | pET15b His SUMO mNeon PPM1H | Addgene in progress |  | human | NEW |
| plasmid | pET15b His SUMO mNeon L66R PPM1H | Addgene in progress |  | human | NEW |
| plasmid | pET15b His SUMO mNeon Δ37 PPM1H | Addgene in progress |  | human | NEW |
| Software, Algorithm | UCSF ChimeraX | PMID:1526424 | RRID:SCR_015872 |  | Reuse |
| Software, Algorithm | Graphpad Prism | Prism 10 version 10.2.3 | RRID: SCR_002798 | www.graphpad.com | Reuse |
| Software, Algorithm | AlphaFold Server | AlphaFold 3 | RRID: SCR_025885 |  | Reuse |
| Protocol | Dephosphorylation of phosphorylated Rab GTPases by PPM phosphatases | [protocols.io](https://www.protocols.io) | dx.doi.org/10.17504/protocols.io.5jyl8d4j7g2w/v1 |  | Reuse |
| Protocol | Expression and purification of PPM1H phosphatase | [protocols.io](https://www.protocols.io) | [dx.doi.org/10.17504/protocols.io.bu7wnzpe](https://doi.org/10.17504/protocols.io.bu7wnzpe) |  | Reuse |
| Protocol | PPM1H Purification from E. coli | [protocols.io](https://www.protocols.io) | [dx.doi.org/10.17504/protocols.io.81wqbnp9vgpk/v1](https://doi.org/10.17504/protocols.io.81wqbnp9vgpk/v1) |  | Reuse |
| Protocol | Preparation of unilamellar liposomes | [protocols.io](https://www.protocols.io) | [dx.doi.org/10.17504/protocols.io.5qpvo3ke7v4o/v1](https://doi.org/10.17504/protocols.io.5qpvo3ke7v4o/v1) |  | Reuse |
| Protocol | Gibson Assembly Cloning | [protocols.io](https://www.protocols.io) | [dx.doi.org/10.17504/protocols.io.eq2lyjwyqlx9/v1](https://doi.org/10.17504/protocols.io.eq2lyjwyqlx9/v1) |  | Reuse |

## Acknowledgements

We are grateful to Dr. Dario Alessi for helpful discussions.

---

**Figure S1.**
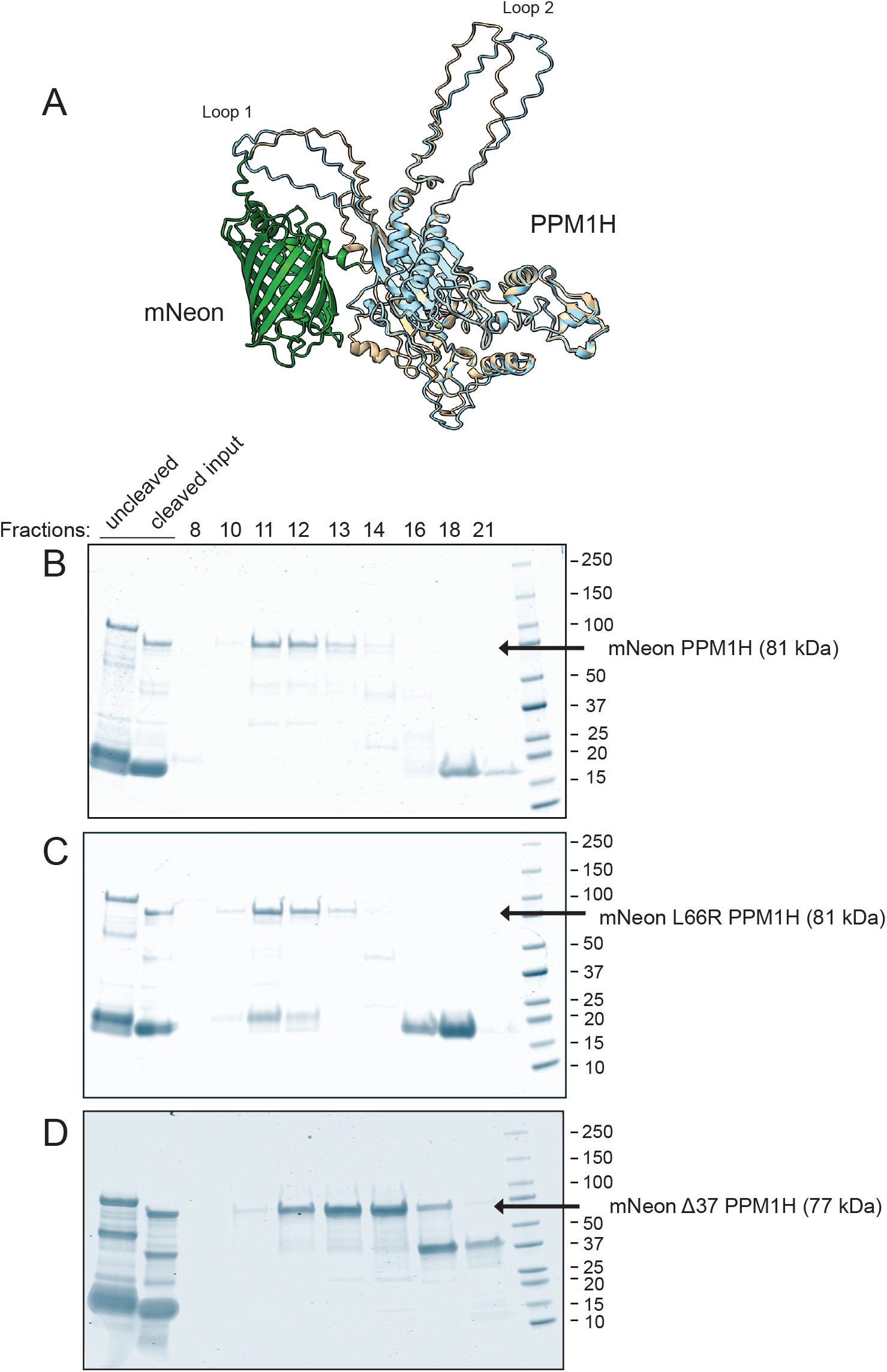
Proteins used in this study. A. Alphafold model of mNeon PPM1H superimposed on wild type PPM1H full length protein. B-D., SDS-PAGE of the gel filtration step of protein purification for (B) mNeon-PPM1H, (C) mNeon L66R PPM1H and (D) mNeon Δ37 PPM1H. Numbers at right indicate molecular weight marker mobility shown in kDa. Also shown are the preparations before and after cleavage to remove the His-Sumo tags.

**Figure S2.**
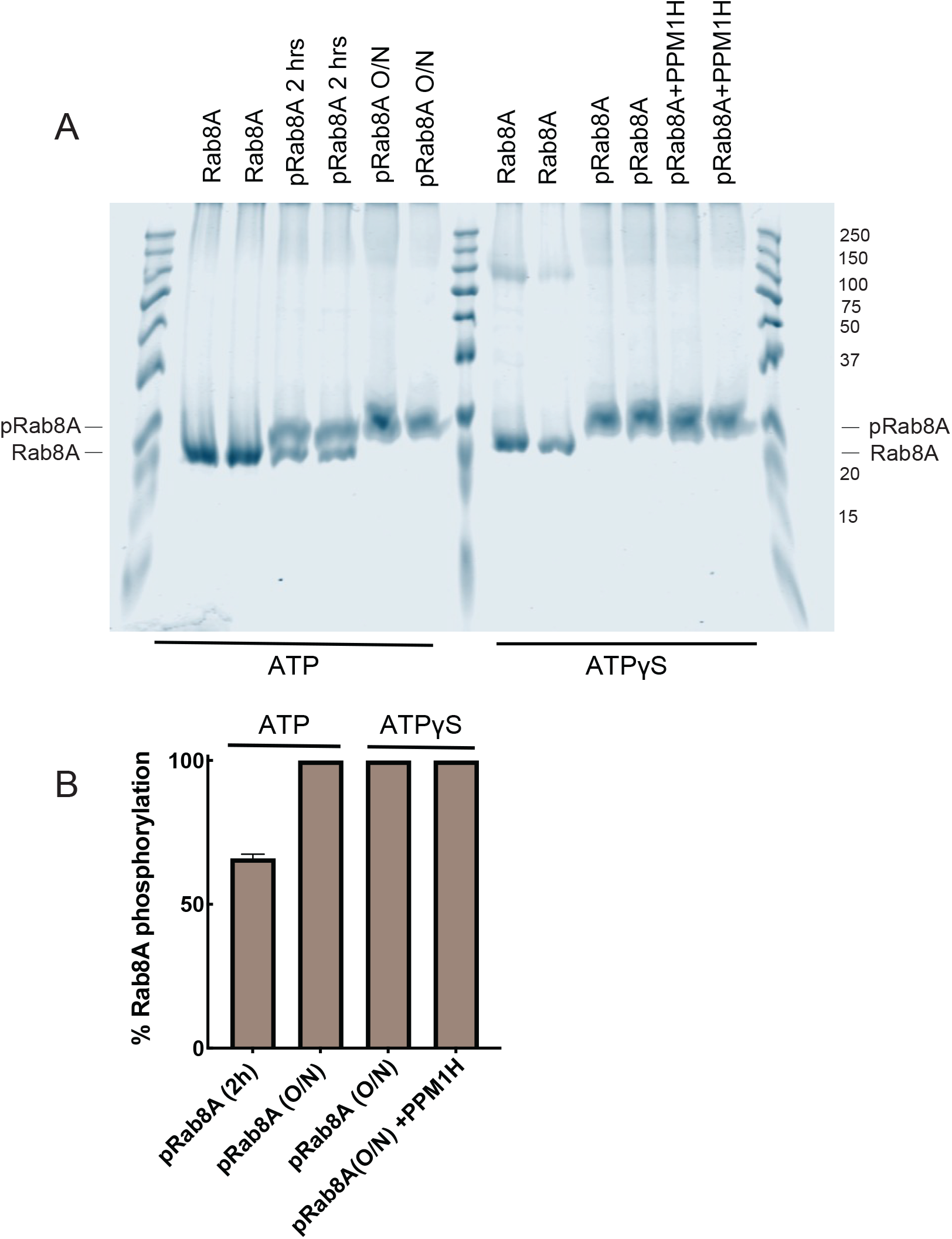
PhosTag gel electrophoresis to monitor MST phosphorylation of Rab8A with ATP (A, left half) or ATP*γ*S (A, right half) as indicated. ATP Reactions were carried out for either 2h or overnight as indicated; ATP*γ*S reactions were carried out overnight; two samples were further treated with PPM1H for 15 minutes at 30°C to check for thiophosphate stability to phosphatase action. B, Quantitation of the gel in A. PhosTag gels were purchased from Fuji Film Wako pure Chemical Corporation, Osaka (195-17991); samples were loaded in sample buffer containing LDS instead of SDS and 1mM ZnCl2, at 120V for 2hr.

